# HuR-Dependent Cooperative Export of miRNAs From Activated Macrophages

**DOI:** 10.1101/2025.08.04.668574

**Authors:** Syamantak Ghosh, Kamalika Mukherjee, Suvendra N. Bhattacharyya

**Affiliations:** RNA Biology Research Laboratory, Molecular Genetics Division, CSIR-Indian Institute of Chemical Biology, Kolkata 700032, India; Department of Anesthesiology, University of Nebraska Medical Center, Omaha, NE 68198-4455, USA; Department of Pharmacology and Experimental Neuroscience, University of Nebraska Medical Center, Omaha, NE 68198-5880, USA

**Keywords:** miRNA, HuR, Ago2, miRNA export, miRNA-HuR interaction

## Abstract

miRNA export is a regulated process vital for maintaining balanced miRNA levels and target gene expression in metazoan cells. RNA-interacting proteins finely adjust the selectivity and specificity of miRNA export, ensuring context-dependent optimal extracellular export of gene-repressive miRNAs in higher eukaryotes. Our findings reveal that activated macrophage cells export miRNAs in a cooperative manner, where elevated expression of miR-122 can significantly enhance the export of miR-146a and selective others. We predict and observe that the cooperative export of repressive miRNAs achieves synchronized upregulation of pro-inflammatory targets in activated macrophage cells. Exploring the molecular basis of this event, we have noted the co-entry of miRNAs into endosomes for their extracellular export, highlighting the selective and cooperative nature of endosome targeting of miRNAs. HuR, a miRNA-binding protein crucial for selectively exporting specific sets of miRNAs from various cell types, facilitates cooperative miRNA entry into endosomes both in the *in vivo* and *in vitro* assay conditions. A catalytic amount of high-affinity miR-122 enhances HuR’s binding to low-affinity miR-146a, facilitating the efficient endosomal compartmentalization of both miRNAs for their co-export via extracellular vesicles (EVs).

## Introduction

MicroRNAs (miRNAs), the 22-nucleotide-long non-coding RNAs, play a pivotal role in the precise regulation of genes, significantly influencing temporal and spatial changes in gene expression at the post-transcriptional level(1, 2). By forming imperfect base pairs with target mRNA sequences, miRNAs effectively induce translational repression and facilitate the degradation of these repressed messages(3). Consequently, regulating miRNA activity is vital for achieving optimal expression of target genes.

Extracellular export is critical for downregulating the functional miRNA pool within eukaryotic cells(4, 5). Recent discoveries have shown that extracellular vesicles (EVs) primarily carry RNA and protein cargo across cellular boundaries(6). The selective export of miRNAs through EVs is influenced by various intrinsic and extrinsic factors, which, in turn, modulate miRNA activity in higher eukaryotes, carrying significant implications for cellular physiology and the onset of diseases(7).

Interactions with specific RNA-binding proteins, including the La antigen, YBX1, and HuR, dictate the export of miRNAs via EVs to neighboring cells (4, 5, 8–13). These proteins selectively enhance the export of miRNAs, thereby regulating the cytosolic miRNA content. In addition to HuR and YBX1, many other RNA-binding proteins have been identified that oversee the miRNA export process across various mammalian cell types. An innovative *in vitro* assay system has been recently developed by our groups to study miRNA entry into endosomes(14). This system has proven invaluable in understanding the selective entry process of miRNAs into endosomes, which is a prerequisite for miRNA export in mammalian cells via EVs.

Observations indicated the export of miR-122 from hepatic cells upon exposure to elevated lipid levels, resulting in inflammation in resident tissue macrophages, particularly in Kupffer cells in murine liver (15, 16). Notably, Kupffer cells initiate the expression of inflammatory cytokines upon receiving the EVs from hepatic cells with miR-122 as cargo(16). Furthermore, LPS-mediated activation of macrophage cells also leads to the secretion of specific miRNAs required for macrophage activation. It is also noteworthy that restricted miRNA export is linked to impaired macrophage activation in LPS-treated cells (17).

Based on the hypothesis that miR-122 presence in macrophages should induce cytokine expression caused by enhanced export of specific cytokine-repressive miRNAs, the EV-mediated export upregulation of miR-146a was documented in macrophages expressing miR-122. The export of miR-146a was found to be essential for miR-122-induced macrophage activation. Observations suggest that miR-146a is cooperatively packaged into endosomes, a process influenced by miR-122 in both *in vivo* and *in vitro* contexts. It was observed that HuR, a strong binder of miR-122, can bind to a low-affinity substrate, miR-146a, in the presence of a “catalytic” amount of miR-122. This observation supports the notion of a cooperative binding mechanism of HuR with miRNAs, suggesting cooperation in miRNA export within activated macrophage cells driven by HuR-mediated entry of miRNAs into endosomes.

In summary, during the early phase of macrophage activation, the cooperative interaction between the stress response protein HuR and miRNAs promotes an increased export of specific miRNAs. This process is initiated by the presence of one or more “catalytic” miRNAs with a strong affinity for HuR to increase the export of secondary miRNAs that are otherwise poor binders of HuR due to their low affinity. These cooperative interactions facilitate endosomal entry and simultaneous export of these secondary miRNA groups, which are crucial for activating and expressing inflammatory cytokines in mammalian macrophage cells.

## Results

### miR-122 induces export of miR-146a from RAW264.7 macrophage cells

Our previous work documented that amino acid starvation-related stress or high lipid exposure induces the secretion of EVs containing miR-122 from hepatic cells(16, 18). This can lead to upregulated cytokine expression in miR-122-EV-recipient macrophages and tissue-resident Kupffer cells, activating them to produce inflammatory cytokines in the high-lipid-exposed murine liver(16). The activation of macrophages is also synchronized with the export of specific miRNAs, a process regulated by the stress response miRNA-binding protein HuR(17).

We anticipate that hepatic miR-122- containing EVs will induce the activation and release of EVs containing specific miRNAs by recipient macrophages (**Fig. 1 A**). Is the internalized miR-122 also part of the miRNA cargo released by recipient RAW264.7 macrophages upon LPS exposure? Or is it involved in EV release by LPS-activated macrophages? After the treatment with LPS for 4 hours, we documented increased levels of miRNAs in EVs secreted by the miR-122- containing EV-treated RAW 264. 7 cells. The increase in miR-146a and miR-122 export by the 4-hour LPS-treated hepatic cell-derived EV-exposed RAW 264.7 cells was consistent with decreases in cellular miR-146a and miR-122 content. In contrast, the cellular levels of let-7a did not show any significant change, although there was an increased association of let-7a with EVs upon LPS treatment (**Fig. 1 B-C**). The export of miRNA correlates with the increased production of inflammatory cytokines IL-1 β and TNF-α in LPS-treated cells, indicating the active status of RAW264.7 cells following LPS exposure (**Fig. 1 D**). Cellular levels of miR-146a have previously been observed to be downregulated during the initial phase of macrophage activation, followed by an elevation at 24 hours of LPS exposure(19). We propose that the export of miR-146a from activated macrophage cells accounts for the enhanced expression of inflammatory cytokines in LPS-treated cells. We also posit that the miR-146a export is regulated by another miRNA, whose presence in activated macrophages causes the upregulation in EV-mediated miR-146a export, which is otherwise not a suitable substrate for the HuR-mediated miRNA export process(20). Given that miR-122 promotes inflammation and miRNAs export in macrophages, we suspected that the EV-derived hepatic miR-122 could enhance the export of miR-146a alongside sets of other miRNAs by activating macrophage cells to drive synchronized upregulation of inflammatory cytokines coupled with export of specific and miR-122 responsive miRNAs.

**Figure 1.**
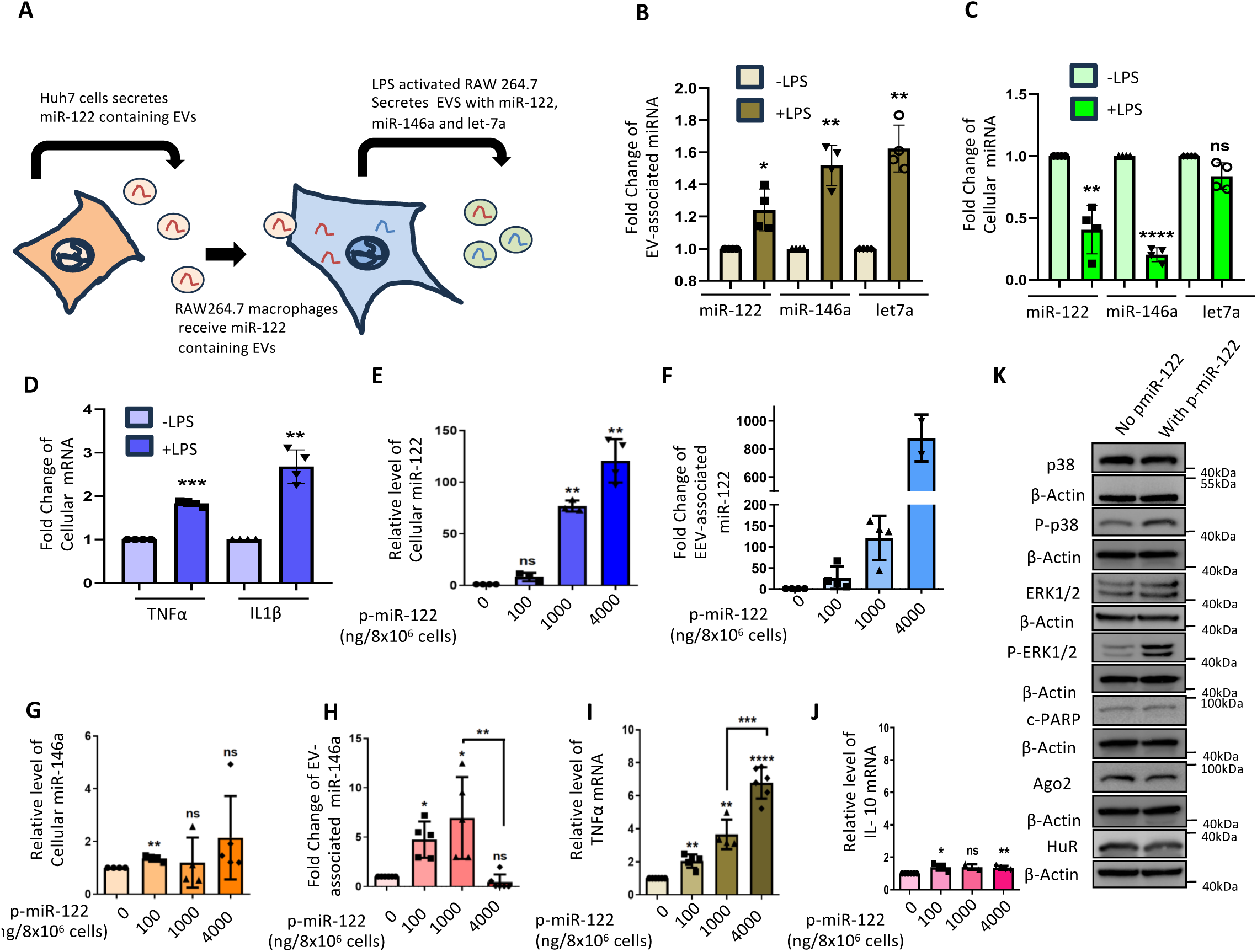
Export of miR-146a from RAW264.7 cells is controlled by miR-122. **A.** Export of miRNAs from macrophage cells received the hepatic miRNAs via EV-mediated delivery. EVs from Huh7 cells, where miR-122 is the predominant cargo packaged in the hepatic EVs, were used to treat naïve RAW264.7 cells for 24 hours and then were activated by LPS (10 ng/ml) for 4 hours. EVs were isolated from-treated RAW264.7 cells. Cellular and EV miRNA levels and cytokine mRNAs isolated from the cells were analyzed. **B-C.** Levels of EV-associated (B), and cellular (C) miRNAs from Huh7 EV-pre-treated and LPS-activated RAW264.7 cells and secreted EVs. (for B panel; n=4, p= 0.0341, 0.0036, 0.0034; for C panle: n=4, p= 0.0088, <0.0001, 0.0575). **D.** Levels of cellular cytokine mRNAs TNFα and IL-1β from LPS-activated RAW264.7 cells, pre-treated with hepatic cell-derived EVs (n= 4, p= 0.0002, 0.0031). **E-F.** Levels of cellular (E), and secreted EV-associated (F) miR-122 in RAW264.7 cells transfected with increasing concentration of pmiR-122 expression vector plasmid. Total RNA from cellular lysate was analyzed to quantify the level of miR-122 by qRT-PCR, and the dataset without the p-miR-122 expression was taken as a unit (n≥3, p= 0.0984, 0.0017, 0.0015). **G-H.** A cooperative effect of miR-146a packaging occurs during EV-mediated export. Cellular (G) and secreted EV-associated (H) miR-146a levels in cells transfected with increasing pmiR-122 expression vector plasmid were measured. Total RNA from cellular lysate was analyzed to quantify the level of miR-146a by qRT-PCR, and the dataset without the p-miR-122 expression was taken as a unit (for G, n≥4, p= 0.0095, 0.2077, 0.7036; For H; n≥5, p= 0.0105, 0.0328, 0.2729 and unpaired t-test p= 0.0022). **I-J.** Cooperative miR-146a export affects the cellular level of cytokine mRNAs. The level of TNF-α mRNA showed an increase with cellular miR-122 level as quantified by qRT-PCR (n≥4, p= 0.0045, 0.0096, <0.0001 and unpaired t-test p= 0.0008) (I). Levels of IL-10 mRNA were also analyzed from the same set of samples (n≥3, p= 0.0324, 0.0636, 0.0089) (J). The dataset without the pmiR-122 expression was taken as a unit to analyze the cytokine mRNAs. **K.** miR-122 expression caused upregulation of inflammatory pathways. P38 MAPK and ERK1/2 activation was evident in western blots done with Phopho-p38 and phosphor ERK1/2 specific antibodies and compared by western blots between the pCI-neo transfected control set and the experimental set ectopically expressed pmiR-122 (at the level of 1000ng pmiR-122 plasmid used to transfect 8×10^6^ cells); the condition that showed cooperative miRNA packaging and export. Data information: In all the experimental data, error bars are represented as mean with SD, ns, nonsignificant, *P < 0.05, **P < 0.01, ***P < 0.001, ****P < 0.0001, respectively. P-values were calculated by a two-tailed paired t-test in most of the experiments unless mentioned otherwise. Relative levels of cellular miRNAs were normalized with U6 snRNA level by 2^-ΔΔCt^ method. The fold change of miRNA was calculated by 2^-ΔCt^ method. Positions of molecular weight markers are marked and shown with the respective Western blots. Normalization was done against the total EV content used for RNA isolation to quantify EV-associated miRNAs. Relative levels of cellular mRNAs were normalized with GAPDH mRNA levels by 2^-ΔΔCt^ metho

Can miR-122 directly influence the export of miR-146a from RAW264.7 cells? Following the miR-122 controlled export of miR-146a, we transfected RAW264.7 cells with miR-122 expression plasmids to induce ectopic expression miR-122 in these cells. We documented an increase in its expression alongside increased EV-mediated export of miR-146a derived from miR-122-expressing RAW264.7 cells in a dose-dependent manner proportional to the rise in cellular miR-122 levels (**Fig.1 E-F**). Interestingly, the EV-associated miR-146a also increases with higher miR-122 expression until a decrease in EV-miR-146a levels occurs when cellular miR-122 content reaches a threshold level (**Fig.1 G**). The cellular miR-146a levels changed reciprocally, as expected. However, changes in cellular miR-146a content were modest until the export of miR-146a was reduced at the highest levels of cellular miR-122 induction, accompanied by a sharp increase in cellular miR-146a content (**Fig.1 H**). These data suggest a biphasic effect of miR-122 on miR-146a expression. While low miR-122 levels promoted miR-146a export and lowered cellular miRNA content, high levels of cellular miR-122 acted as a competitive inhibitor of miR-146a export from RAW264.7 cells.

Does miR-122-induced export of miR-146a enhance the expression of inflammatory cytokines in RAW264.7 cells? Consistent with our previous observation of EV-associated miR-122 treatment-induced expression of inflammatory cytokines in macrophages (16, 18), we documented an enhancement of TNF-α in miR-122-expressing cells with a marginal effect on IL-10 anti-inflammatory cytokine expression (**Fig. 1I, J**). Up to a concentration of 1000 ng plasmids used to transfect 8×10^6^ cells, cooperative miR-146a export was observed with a modest level of cellular miR-122. We assessed the status of immune activation in miR-122 expression in those cells. We observed that miR-122 activates the p-p38 MAPK and p-ERK1/2 pathways in miR-122-expressing RAW264.7 cells (**Fig. 1K**), as observed in hepatic macrophages(18).

In LPS-activated macrophages, we observed miR-146a induction occurring during the late phase of activation (24h), which was associated with the miR-146a-dependent biogenesis of “secondary” miRNAs from respective pre-miRNAs. This “cooperative biogenesis” of miRNAs results in increased mature miRNA levels for secondary miRNAs regulated by miR-146a and a concomitant decrease in the respective pre-miRNA levels (21). Interestingly, miR-122 expression, which enhances miR-146a export, does not induce coordinated biogenesis of “secondary” miRNAs, as there are no significant changes in the mature and pre-miRNA levels for “secondary” miR-125b or miR-143-3p (**Fig. S1 and B**), clearly showing the coordinated miRNA export phase is separate to that of coordinated biogenesis reported earlier for the late phase of macrophage activation.

### miR-122 induced miR-146a export augments the inflammatory responses

We assume that the export of miR-146a occurring in 4h LPS-activated or miR-122-expressing RAW264.7 cells is essential for activating macrophage cells. To test whether miR-146a export may be linked to the activation of RAW264.7 cells, we block the export of miRNAs using GW4869 in cells expressing miR-122 in an inducible manner(12). Inducible miR-122-mediated expression of miR-122 in RAW264.7 cells has been achieved with Doxycycline(19). With miR-122 expression, blocking the EV-mediated export resulted in a reduction of TNF-α and IL-1β mRNA levels. To confirm that the effect of blocking miRNA export on IL-1β and TNF-α is due to an increased cellular level of miR-146a, we block the activity of miR-146a by transfecting cells with anti-miR-146a oligomers. Inactivation of miR-146a resulted in partial restoration of expression of both IL-1β and TNF-α (**Fig. 2A-B**). This is consistent with the idea that miR-146a, among the other cooperatively exported miRNAs by miR-122, regulates the expression of the inflammatory cytokines in miR-122-activated macrophages. Thus, accumulated miR-146a inhibition in GW4869-treated cells can only partially rescue the IL-1β and TNF-α expression.

**Figure 2.**
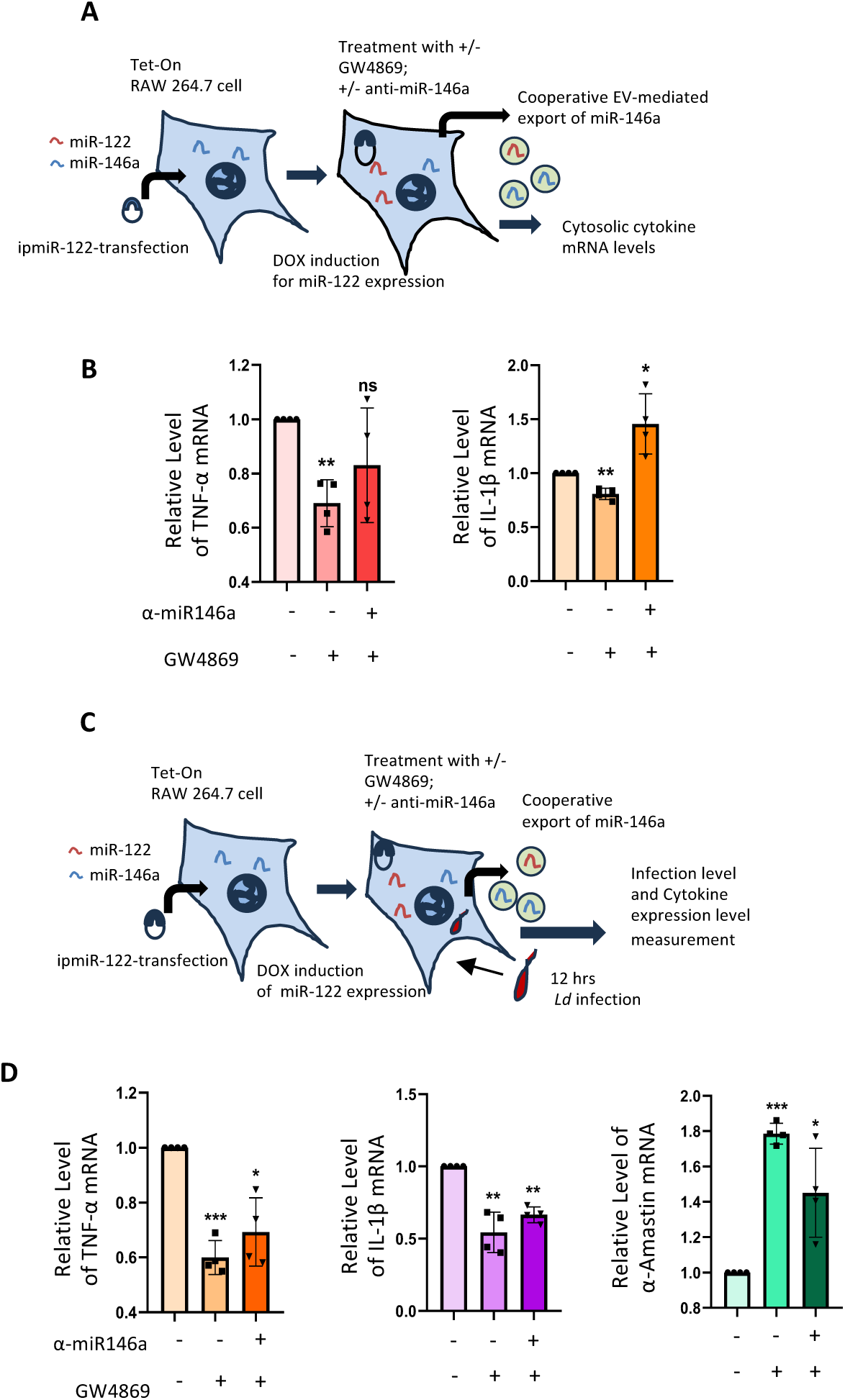
Cooperative export of miR-146a by miR-122 is essential for proinflammatory response. **A.** Scheme of the experiment. RAW264.7 cells stably expressing Tet-On expression cassette were transfected with Doxycycline inducible-miR122 expression vector. These cells were subjected to miR-146a inhibitor (anti-miR-146a oligos, 30 nM, 24 hours) and EV secretion blocker-GW4869 treatment (10 µM, 16 hours) to test their effect on cytokine expression. miR-122 expression was induced with doxycycline (400 ng/ml, 24 hours). The cellular levels of key cytokine mRNAs were measured. **B.** The cellular levels of TNF-α and IL-1β mRNAs under the different treatment conditions were analyzed by RT-PCR (TNFα; n= 4, p= 0.0056, 0.2063; IL-1β; n= 4, p= 0.0053, 0.0470). **C.** Scheme of the experiment done with *Leishmania donovani* infection of RAW264.7 macrophage. Cells stably expressing Tet-On expression cassette were transfected with Doxycycline inducible-miR122 vector (ipmiR-122). These cells were subjected to miR-146a inhibitor (anti-miR-146a oligos, 30 nM, 24 hours) and EV secretion blocker-GW4869 treatment (10 µM, 16 hours). miR-122 expression was induced with doxycycline (400 ng/ml, 24 hours), which was followed by infecting the cells with *Leishmania donovani* for 6 hours. Then, the cells were allowed to recover for another 12 hours post-infection. The cellular levels of a key cytokine mRNA and a pathogen-specific mRNA were measured. **D.** The cellular levels of TNF-α and IL-1β cytokine mRNAs, along with α-Amastin level, were analyzed by qRT-PCR to determine the level of inflammatory condition and *Ld* infection level under the different conditions of treatments and infection. (For TNF-α, n= 4, p= 0.0010, 0.0158; for IL-1β, n= 4, p= 0.0074, 0.0012; for α-Amastin, n= 4, p= 0.0001, 0.0372). Data information: In all the experimental data, error bars are represented as mean with SD, ns, nonsignificant, *P < 0.05, **P < 0.01, ***P < 0.001, ****P < 0.0001, respectively. P-values were calculated by a two-tailed paired t-test in most of the experiments unless mentioned otherwise. Relative levels of mRNAs were normalized with GAPDH mRNA levels by 2^-ΔΔCt^ method.

miR-146a plays a crucial role in the anti-inflammatory response. The upregulation of miR-146a is essential for macrophage infection by the pathogen *Leishmania donovani*. The miR-146a-containing extracellular vesicles (EVs) derived from infected macrophages are also getting transferred to naïve macrophages, polarizing them to M2-like by increasing the miR-146a content in the recipient macrophages, which simultaneously downregulate inflammatory cytokines in recipient cells(20). We documented an enhanced level of infection, measured by the internalization of the pathogen-specific gene α-amastin in cells where GW4869 blocks the export of miR-146a. The experiments were conducted in RAW264.7 expressing miR-122. We observed a reversal in the expression of α-amastin when the cells were pre-transfected with anti-miR-146a oligonucleotides, confirming that the effect of GW4969 on the infection level was primarily due to the increased miR-146a level (**Fig. 2 C-D**). Changes in IL-1β and TNF-α cytokines also followed a similar trend in infected cells where miRNA export was blocked, with partial recovery in expression when treated with anti-miR-146a oligos under comparable conditions (**Fig. 2 C-D**).

### Cooperative entry of miRNA into endosomes is dependent on HuR

How can the presence of one miRNA enhance the export of a secondary miRNA, and how is the cooperative export of miRNAs mechanistically possible? miRNAs are needed to be transported to endosomes for subsequent EV-mediated export(12). We have recently developed an *in vitro* miRNA import assay with isolated endosomes to study the endosomal entry of miRNAs. We documented a concentration-dependent entry of single-stranded miRNAs into the endosomal lumen, which becomes resistant to RNase used to digest the non-internalized miRNA pool after the import reaction(14). Does miRNA entry into endosomes occur cooperatively? To test this, we conducted the *in vitro* import assay for miR-146a in the presence of increasing concentrations of miR-122, using endosomes isolated from C6 glioblastoma cells(14). Interestingly, in line with our *in vivo* observations, we found a concentration-dependent positive effect of miR-122 on miR-146a entry into endosomes (**Fig. 3 A-B**). This data is consistent with the *in vivo* results we obtained with pmiR-122 transfected RAW264.7 cells for miR-146a, where the presence of miR-122 at a low “catalytic” amount positively affected the export of miR-146a.

**Figure 3.**
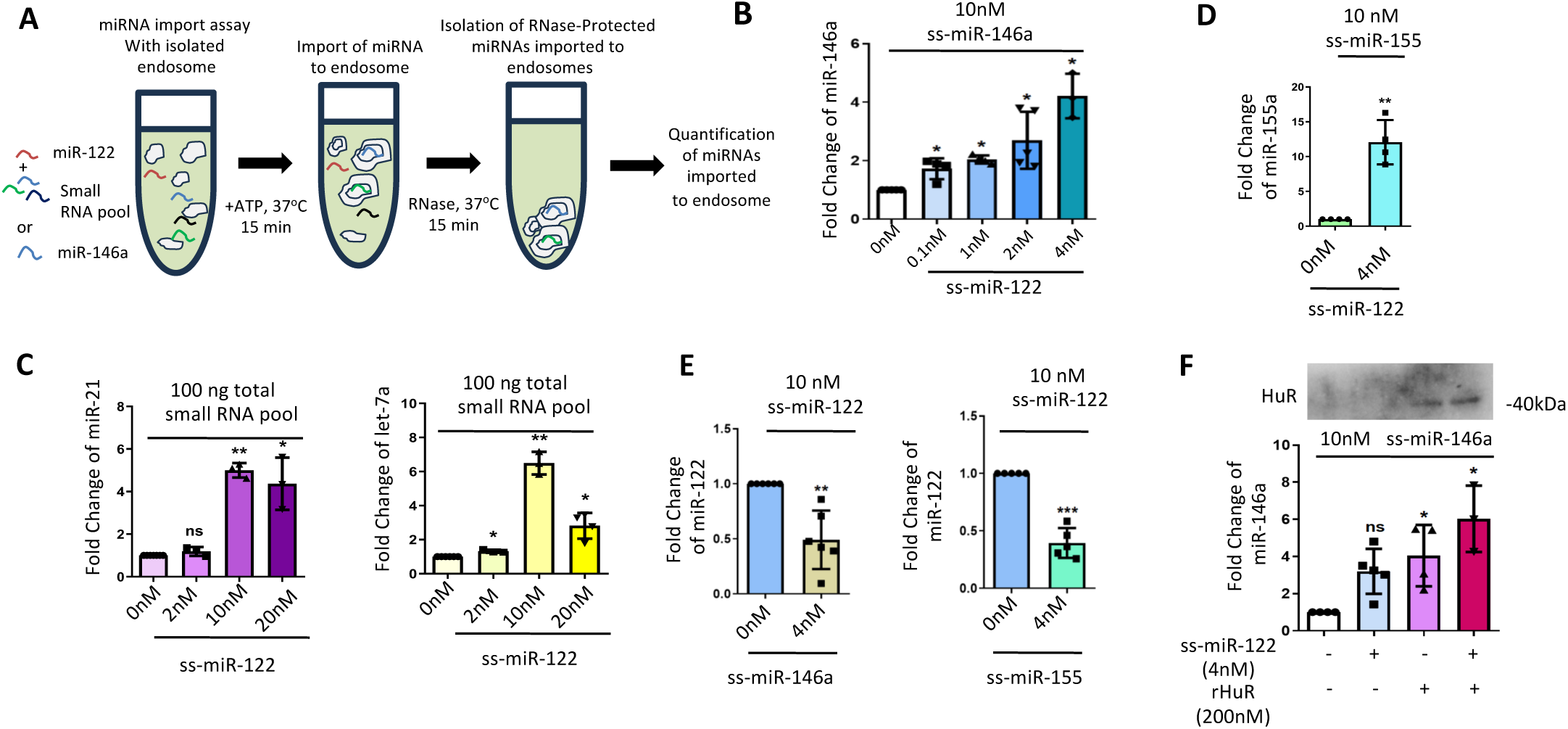
miR-122 promotes cooperative packaging of miRNAs into endosomes. **A.** Scheme of the *in vitro* endosome targeting assay of miRNAs. The graphical diagram represents the cooperative miRNA packaging achieved in an *in vitro* assay performed with enriched endosomes isolated from C6 rat glioma cells. Under different conditions, as required by specific experiments, synthetic single-stranded miRNAs such as ss miR-122, ss miR-146a, ss miR-155, or a pool of small RNAs isolated from RAW264.7 cells were used as cargo miRNA to observe the cooperation and competition between multiple miRNAs for the packaging into endosomes. **B.** Cooperative effect of endosomal packaging of two species of miRNAs. Synthetic ss-miR-146a (10nM) was added as the cargo and incubated for 30 minutes with increasing amounts of ss-miR-122 (0, 0.1, 1, 2, 4nM). Excess non-imported RNA level was minimized by RNase treatment followed by endosome vesicle re-isolation. QRT-PCR quantified the packaged miRNAs. The dataset with 0 nM ss-miR-122 was considered as a unit (n≥3, p= 0.0273, 0.0227, 0.0177, 0.0184). **C.** With endosomal suspension from C6 cells, 1000ng/ml of small RNA pool isolated by miRVana kit (Thermo Scientific) was added with an increasing concentration of synthetic single-stranded (ss) miR-122. miRNA packaging reaction was carried out for 30mins followed by an RNase protection and vesicle re-isolation (details in the Methods section). QRT-PCR measured packaged levels of various miRNAs, and the value of individual small RNA in the presence of 0nM ss-miR-122 was considered a unit. Quantitative measurement for endosome targeted miR-21 (left) and let7a (right) miRNA (miR-21; n=3, p= 0.2671, 0.0024, 0.0414; let-7a; n≥3, p= 0.0239, 0.0049, 0.0168) are shown. **D.** Influence of ss-miR-122 (4nM) on endosomal packaging of 10nM of ss-miR-155a. Excess reactant RNA was minimized by RNase reaction and vesicle re-isolation. The packaged miRNA was quantified by qRT-PCR and the dataset with 0nM ss-miR-122 was considered as a unit (n=4, p= 0.0061). **E.** Effect of ss-miR-122 (10nM) on endosomal packaging was studied in the presence of ss-miR-146a or ss-miR-155 (4nM). After 30 minutes of incubation, excess unpackaged RNA was eliminated by RNase reaction and vesicle re-isolation. The internalized miRNA was quantified by qRT-PCR and the dataset with 0nM ss-miR-146a (left panel) or 0nM ss-miR-155 (right panel) was considered as unit (left panel n=6, p=0.0054; right panel n=5, p=0.0005). **F.** Cumulative cooperative effect of ss-miR-122 (0.1nM) and recombinant HuR (rHuR, 50nM) on endosomal packaging of ss-miR-146a (10nM). The reaction was carried out for 30 minutes; excess unpackaged RNA was depleted by RNase reaction and vesicle re-isolation. QRT-PCR quantified the packaged miRNA, and the dataset without ss-miR-122 and rHuR was considered a unit. Western blot shows the level of HuR from the samples after the vesicles were re-isolated (n≥3, p= 0.0521, 0.0348, 0.0396). Data information: In all the experimental data, error bars are represented as mean with SD, ns, nonsignificant, *P < 0.05, **P < 0.01, ***P < 0.001, respectively. P-values were calculated by a two-tailed paired t-test in most of the experiments unless mentioned otherwise. The fold change of miRNA was calculated by 2^-ΔCt^ method. Endosomal miRNA level-normalization was done against the total endosomal content used for the assay to quantify endosome-associated miRNAs. Positions of molecular weight markers are marked and shown with the respective Western blots.

Does the miR-122 effect remain exclusive to miR-146a? We incubated the small RNA pool isolated from C6 cells and treated it with increasing concentrations of synthetic single-stranded (ss) miR-122. We observed a dose-dependent increase in the import of let-7a and miR-21 by miR-122. Interestingly, while the 10mM concentration of miR-122 enhances the entry of miR-21 and let-7a into endosomes, a 20mM concentration of miR-122 substantially reduces the import level of let-7a and begins to show a decrease in miR-21 endosomal import levels as well.

In contrast, at higher concentrations, miR-122 exhibited an inhibitory competitive impact on miR-146a (**Fig. 3C and Fig.1 G**). When synthetic miR-155 was used as a secondary substrate at a 4nM miR-122 concentration, cooperative entry for miR-155 was also observed in an endosomal targeting assay *in vitro* (**Fig.3 D**). Interestingly, with the reversal of the roles of miR-146a or miR-155 used at lower concentrations, both miRNAs inhibited the endosomal entry of synthetic ss miR-122 (**Fig. 3E**). This data suggests that the cooperative effect of one miRNA on another is unidirectional and specific. For cooperative effect, under varying concentrations, the same miRNAs may act as competitive inhibitors when used at higher concentrations. In contrast, at “catalytic” concentrations, the cooperativity of miRNA entry into endosomes, driven by a primary catalytic miRNA such as miR-122, for subsequent EV-mediated export of a set of co-regulated miRNAs has been documented.

HuR is known to regulate the miRNA export process by reversibly binding the miRNAs to facilitate the endosomal miRNA entry(12, 14). To test the effect of HuR on the cooperative entry of miRNAs into endosomes, we conducted an endosome import assay for miR-146a in the presence of HuR and miR-122, documenting enhanced endosomal import of miR-146a when recombinant HuR and miR-122 are added separately. The import of miR-146a in the presence of miR-122 is significantly improved with recombinant HuR (**Fig. 3 F**), suggesting an additive effect of miR-122 and HuR on miR-146a entry into endosomes.

To confirm that endosomal compartmentalization of miRNAs occurs cooperatively in the *in vivo* context and that EV-associated miRNA export is proportional to this cooperative endosomal targeting, we expressed miR-122 in RAW264.7 cells alone or with HA-HuR to evaluate the effects on subcellular miR-122 and miR-146a content, as well as changes in their EV association. With the expression of miR-122, there was an increase in EV content released by pmiR-122-expressing cells, which was significantly enhanced with the co-expression of HA-HuR in RAW264.7 cells (**Fig.S2 A-D**). This may explain the activation state of RAW264.7 cells, which is associated miR-122 expression and improved miRNA and EV export. We performed subcellular fractionation of cell lysates on an Optiprep-density gradient to separate endosomes and the ER, and we quantified the amounts of ER and endosome-associated miRNAs obtained from pCI-neo control, pmiR-122, and pmiR-122 plus pHA-HuR co-transfected cells (**Fig. S3A-B**)(14, 22). With the transfection of pmiR-122 and pHA-HuR, we documented an increased presence of EV- and endosome-associated miR-122 (**Fig. S3C**). At the same time, a clear enrichment of miR-146a content in EVs and endosomes was also noted (**Fig. S2 D**). Thus, both *in vitro* and *in vivo*, miR-122 and HA-HuR positively influence the endosomal targeting of miR-146a cooperatively. This explains the enhanced miR-146a export via EVs from cells expressing low levels of miR-122.

### High-affinity miR-122 enhances the binding of HuR with low-affinity substrate miR-146a

How does the cooperative entry of miRNAs into endosomes occur *in vitro and in vivo*? The stress response protein HuR is a miRNA binder and is necessary and sufficient for miR-122’s import into endosomes and EV-mediated export in hepatic cells under stress(12). HuR is also required for exporting miRNAs like let-7a and miR-155 from LPS-activated RAW264.7 macrophage cells(17). These miRNAs are found to be regulated cooperatively by miR-122a (**Fig. 3C and D**), while cooperative entry of miR-146a into endosomes is also facilitated by HuR (**Fig.3F**).

How HuR promotes the cooperative entry of miRNAs into endosomes remains to be seen. HuR binds with its miRNA substrates with differential affinity, possibly determined by the presence or absence of AU-rich sequences on the RNA substrates. We used HA-HuR immobilized on Protein-G agarose beads, which were incubated with a mix of ss miR-122 and ss miR-146a. To identify the cooperative effect of miR-122 on miR-146a HuR binding, we incubated 10nM miR-122 with HuR immobilized beads in the presence of increasing concentrations of miR-146a. For a fixed amount of miR-122 as substrate, the increasing concentration of miR-146a caused a concentration-dependent decrease in miR-122 binding to HuR. Conversely, miR-146a binding remained low until a threshold concentration of miR-146a caused a jump in HuR-miR-146a binding at 4nM (**Fig. 4A-B**). With a fixed amount of miR-146a used in a binding assay, increasing the miR-122 concentration resulted in a sharp rise in miR-146a binding with 1 and 2 nM of miR-122 present, while a drop in miR-146a binding to HuR was noted with 4nM of miR-122 (**Fig. 4 C).** This data suggests that the cooperative binding of miR-146a to HuR occurs in the presence of lower concentrations of miR-122. The cooperativity of miRNA binding with HuR can explain the cooperative miRNA export we have observed in macrophages. We posit that in the presence of a catalytic amount of miR-122 (the high-affinity binder), the HuR-miR-122 complex forms, allowing the formation of the miR-146a-HuR complex (**Fig. 4 D**). HuR forms oligomers with the RNA substrates. HuR oligomerization on AU-rich sequences of target mRNAs is required for miRNA replacement from target mRNAs and miRNA derepression(23). Thus, HuR oligomerization in the presence of the high-affinity substrate miR-122 may increase the affinity of oligomeric HuR for the low-affinity substrate miR-146a as the later get trapped in a co-oliogomers of HuR and miR-122 (**Fig.4D**).

**Figure 4.**
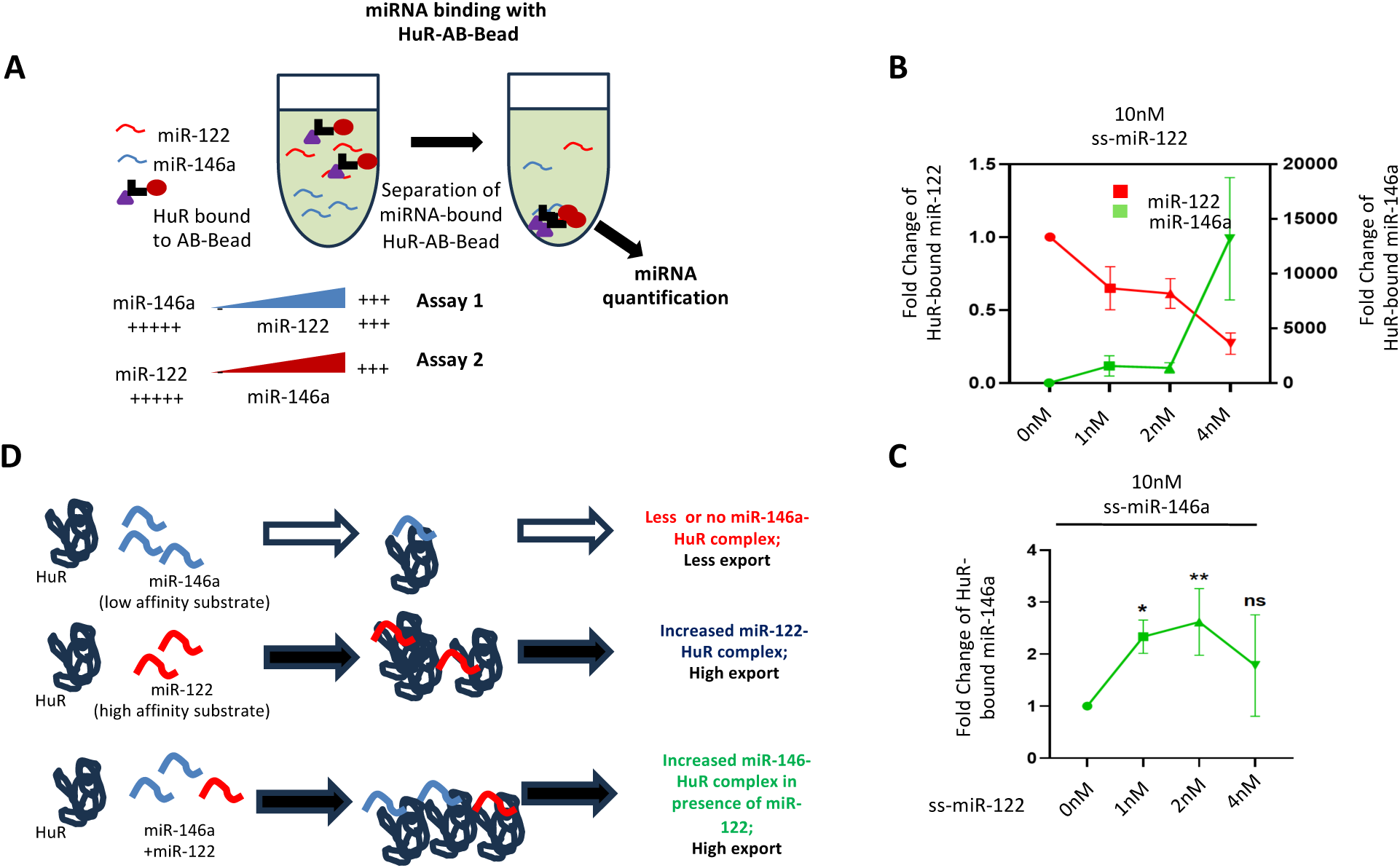
Cooperative binding of low-affinity miR-146a to HuR in the presence of high-affinity miR-122. **A.** Scheme of the experiments. The HA-HuR expression vector was transfected in HEK293 cells, and from the cellular lysate, HA-HuR was immunoprecipitated and immobilized on HA-antibody-bound Protein-G agarose beads. Equal amounts of immobilized HA-HuR were incubated in a HuR assay buffer (details in the Methods section) for 15 minutes at room temperature. After the reaction, beads were pelleted down and washed to remove unbound components. The beads were divided equally to analyze miRNAs and proteins from these samples. **B.** Suggested as assay 2 in panel A; synthetic ss-miR-122 (10nM) was added in the immobilized HA-HuR reaction system along with varying amounts of synthetic ss-miR-146a (0, 1, 2, 4nM). After the incubation, beads were thoroughly washed to remove excess or non-specific RNAs, and then samples were divided equally for RNA and protein analysis. The HuR-associated miR-122 and miR-146a levels were analyzed by qRT-PCR considering the dataset of 0nM ss-miR-146a as a unit (n=2). The amount of HuR isolated as immunoprecipitate was used for normalization. **C.** Suggested as assay 1 in panel A; synthetic ss-miR-146a (10nM) was added in the immobilized HA-HuR reaction system along with varying amounts of secondary miRNA species synthetic ss-miR-122 (0, 1, 2, 4nM) and HuR association reaction was carried out. After the reaction, beads were thoroughly washed to remove excess reactants, and then samples were divided equally for RNA and protein analysis. QRT-PCR analyzed the HuR-associated miR-146a levels by considering the dataset of 0nM ss-miR-122 as a unit (n=3, p= 0.0186, 0.0096, 0.0701). The amount of HuR isolated as immunoprecipitate was used for normalization. **D.** Possible model of cooperative binding of HuR with miR-146a in the presence of miR-122. High affinity miR-122 promotes the low binder miR-146a binding with HuR, where HuR oligomerization favored by miR-122 may assist the miR-146a binding or entrapment within HuR-oligomers. Data information: In all the experimental data, error bars are represented as mean with SD, ns, nonsignificant, *P < 0.05, **P < 0.01, ***P < 0.001, respectively. P-values were calculated by a two-tailed paired t-test in most of the experiments unless mentioned otherwise. The fold change of miRNA was calculated by 2^-ΔCt^ method.

## Discussion

Our previous observations suggest that LPS induces the uncoupling of miRNAs from phosphorylated Ago2, which leads to the accumulation of Ago2 free from miRNA(24). The fate of Ago-free miRNAs has not been investigated previously but was expected to be exported out from activated macrophages(17, 20). The coordinated export may indicate how the balanced expression of Ago-uncoupled miRNAs can serve as substrates for EV-mediated export. However, the relationship between the cooperative export of miRNAs and Ago2’s miRNA unbinding remains unknown.

The cooperative import of miRNAs into endosomes is linked to the co-export of miRNAs documented in RAW 264.7 cells. The miRNA-binding protein HuR is essential for the cooperative import of miRNAs into endosomes and the EV-mediated co-export of miRNAs. HuR’s binding with high-affinity miR-122 enhances the binding of low-affinity miRNA substrates to HuR, facilitating the co-entry of both miRNAs into endosomes. We documented earlier that the RRMIII of HuR is crucial for miRNA export and miRNA-binding activity(12). At the same time, the hinge region of HuR is responsible for HuR ubiquitination and EV-mediated miRNA export(12). We posit that HuR’s binding with high-affinity miRNAs may induce a conformational change that enhances the binding with secondary low-affinity miRNAs, possibly through other two RRMs of HuR, or it may be due to accelerated oligomerization promoting the entrapment of Ago2-free miRNAs with the oligomeric form of HuR(25). The HuR-miRNA complex subsequently activates RalA GTPase and causes selective but cooperative import of miRNAs into the endosomes(14). The HuR-miRNA complex is also known to transfer the miRNAs to the syntaxin protein STX5, which acts synergistically with HuR for endosome targeting and EV-mediated miRNA export(22). It is unclear how STX5 or RalA may affect the cooperativity of miRNA import into endosomes(22).

The coordinated miRNA regulation and export observed in the early phase of macrophage activation ensure the systematic inactivation and export of miRNAs from macrophage cells. This allows the cells to adjust to activation signals and simultaneously permits the coordinated expression of cytokine-encoding miRNA-repressed mRNAs in activated macrophages. Thus, the EV-mediated coordinated export, in turn, regulates specific sets of miRNAs and their target genes simultaneously.

During the late phase of macrophage activation, coordinated re-repression of miRNA targets must be ensured to prevent damage from unregulated and excess cytokine production detrimental to the macrophage(21). The coordinated biogenesis of secondary miRNAs is achieved by increased expression of primary miRNAs, such as miR-146a, which induces the biogenesis of sets of secondary miRNAs(19). miR-146a in RAW264.7 cells shows a biphasic expression pattern. The cooperative export-mediated downregulation of miR-146a at the early phase of activation by miR-122 or LPS causes a surge in cytokine expression. However, with increasing induction of miR-122, it gradually acts competitively to block the export of miR-146a, allowing the accumulation of miR-146a, possibly resulting in coordinated biogenesis of secondary miRNAs occurring during the late phase of macrophage activation, as it also noted in LPS-treated macrophages (**Fig. 5 A-B**). Therefore, miR-146a is one of the key regulatory miRNAs controlled by coordinated export that ensures regulated cytokine production. At a later stage, the accumulation of miR-146a drives coordinated miRNA biogenesis to suppress many cytokine expressions during the late activation phase. The importance of both the coordinated regulation of miR-146a export and miR-146a-driven miRNA biogenesis in mammalian macrophage cells have clear physiological implications in inflammatory responses and tissue-specific inflammation regulation in the liver under Non-alcoholic steatohepatitis (NASH) condition or in degenerating brains on exposure to amyloid proteins (**Fig. 5C-D**).

**Figure 5.**
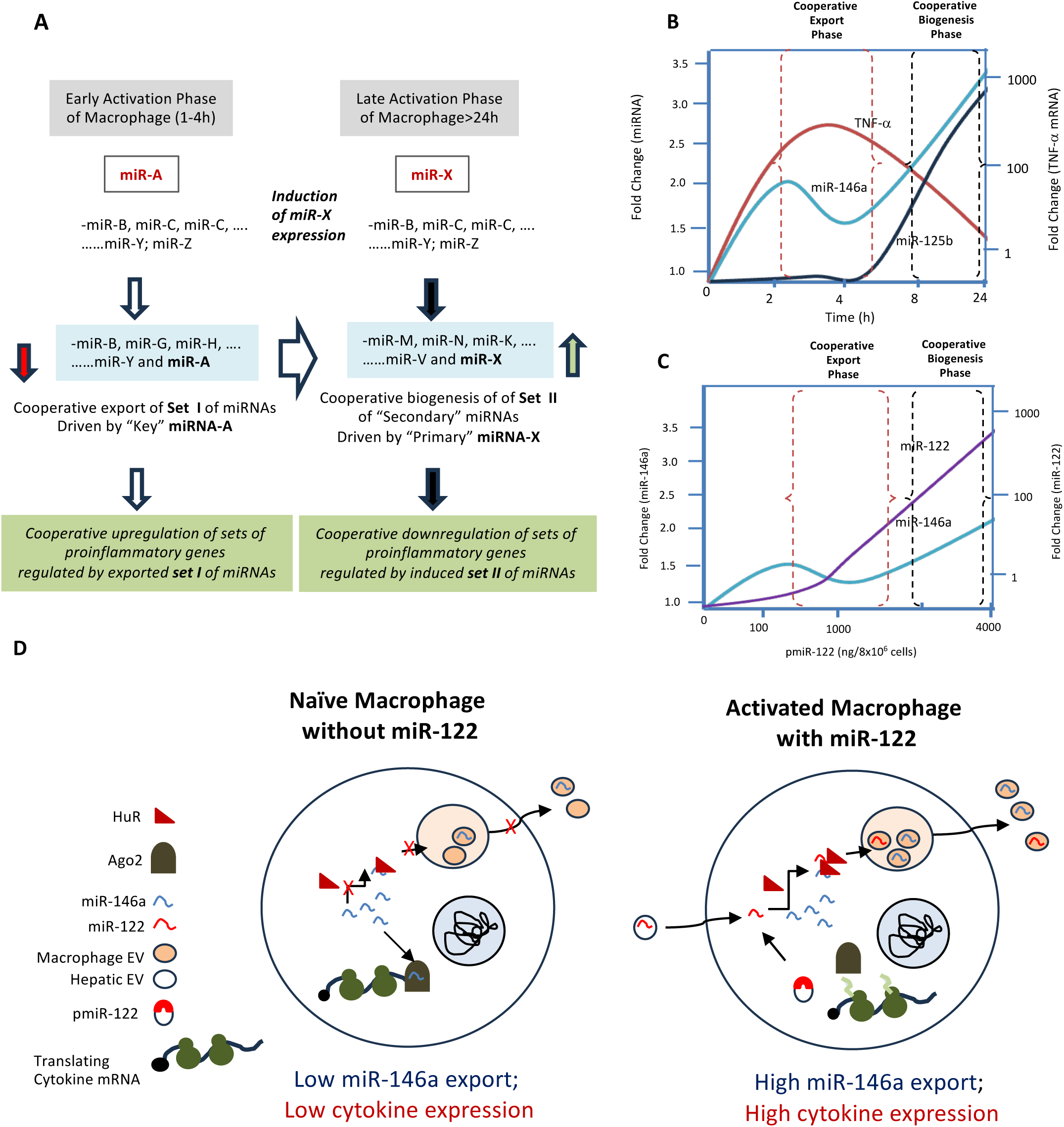
Biphasic cooperative regulation of miRNAs in mammalian macrophages. **A.** The biphasic regulation of miRNAs is observed in mammalian cells. Induction of cooperative export induced by exogenous miR-122 or specific endogenous miRNA (miR-A) caused cooperative export of secondary miRNAs regulating specific cytokine and related mRNAs to ensure robust and cooperative expression during the early phase of macrophage activation, essential for pro-inflammatory cytokine expression. The late anti-inflammatory reactions caused by a specific member of the miRNAs induced by late-phase associated miRNA(s) (miR-146a; miR-X) that causes coordinated biogenesis of miRNAs to ensure repression of cytokine expression in the late phase or overly activated macrophage to protect the macrophage from apoptotic death; as part of the essential LPS-tolerance mechanism. **B.** The available data suggest biphasic expression level changes for miR-146a in activated macrophages(19). The secondary miR-143-5p follows a late response expression pattern that accounts for the bell-shaped TNF-α expression pattern. This data shows how cooperative export and biogenesis contribute to immune activation regulation in macrophage cells. **C.** Dose-response data of miR-122 and miR-146a suggest that miR-122, at low concentration, contributes to the cooperative export of miR-146a while, at higher concentration, competitively prevents the export of miR-146a. Thus, miR-122 contributes to the cooperative biogenesis step of miRNAs induced by accumulated miR-146a to optimize immune response. **D.** A graphical summary of the macrophage activation process controlled by cooperative export of miR-146a by miR-122. Expression or uptake of miR-122 into macrophages resulted in induced export of miR-146a, repressive miRNAs for the expression of inflammatory cytokines, which resulted in macrophage activation.

We have identified miR-122 as the central hepatic cell-derived miRNA responsible for inflammation in murine livers exposed to a high-fat diet. The miR-122-containing EVs released by hepatic cells upon exposure to high lipids are taken up by Kupffer cells, where they induce inflammatory cytokines. We posit that miR-122, upon entering the macrophage, induces the coordinated export of anti-inflammatory miR-146a as a feedback mechanism to deliver it to hepatic cells to reduce miR-122 expression there. The reduction in miR-122 expression by miR-146a and miR-146a-containing EVs has been documented in other contexts, supporting our hypothesis of reciprocal regulation occurring in fatty liver (20). Thus, macrophage-hepatocyte crosstalk via EVs, with coordinated miRNA export, may be essential in controlling metaflammation-related liver damage.

## Experimental Procedures

### Cell culture and cell transfections

HEK293 and C6 glioblastoma cells were cultured in Dulbecco’s Modified Eagle’s medium (DMEM; Gibco) supplemented with 10% heat-inactivated fetal calf serum (HI-FCS, Gibco) and 1% of Penstrep (Gibco). RAW264.7 murine macrophage cells were grown in RPMI1640 medium (Gibco) with 2mM L-glutamine and 0.5% β-mercaptoethanol along with 10% FBS and 1% Penstrep. These cells were grown in a 37°C incubator with 5% carbon dioxide. RAW264.7 cells were stimulated with LPS (10 ng/ml for 4 hours) for macrophage activation from Escherichia coli O111:B4 (Calbiochem). For EV-free growth medium, 10% EV-depleted FBS (made by ultracentrifugation of FBS at 1,000,00g for 4 h) was added to DMEM or RPMI-1640 for respective cell types. For HEK293 and C6 cells, all transfections of plasmids and co-transfection with miRNA inhibitor (anti-miR oligonucleotides, 30 nM) were performed using Lipofectamine 2000 reagent (Life technologies) according to the manufacturer’s protocol.

### Extracellular Vesicles isolation and characterization

Huh7, and RAW264.7 cells transfected with specific plasmids were split 24h post-transfection into 90mm plates and incubated for 24 hours to achieve 70-80% confluency. The conditioned media were subjected to EV isolation as described in a previous report, with minor modifications (26). Cells were grown in growth medium supplemented with EV-depleted serum to prevent any interference from exosomes or EVs present in serum. The conditioned media was clarified for cellular debris and other contaminants by centrifugation at 2,000xg for 10 minutes and 10,000xg for 30 minutes followed by filtering through a 0.22-µm filter for further clearing. EVs were isolated by ultracentrifuging the cleared conditioned media at 100,000xg for 90 min. When analysis of proteins was required, an additional ultracentrifugation of the cleared conditioned media was done on 30% sucrose cushion at 100,000g for 90 minutes, and the layer of EVs was further diluted with 1X phosphate buffered saline (PBS) by ultracentrifugation at 100,000xg for 90 minutes for obtaining EV pellet.

For characterization, the EVs were diluted 10 fold with PBS and 1 ml of these diluted EV samples were injected in the Nano-particle tracker (Nanosight NS-300) and the number, size, and other parameters of EVs were measured.

### EV treatment of recipient cells

After EV isolation, the pellet was resuspended in fresh RPMI-1640 supplemented with 10% EV– depleted FBS and 1% Pen-Strep followed by filtration through a 0.22-μm filter unit. The EVs were then added to the recipient macrophage cell line for 24 h. Post-LPS treatment was given at a dose of 10 ng/ml for 4 h after 20 h of EV treatment to macrophages. For treatment, EVs isolated from 15×106 cells (5X 60 mm plate) were added to 2.4×106 recipient cells (1X 60 mm plate).

### Parasite culture and infection to macrophage cell line

*Leishmania donovani* (Ld) strain AG83 (MAOM/IN/83/AG83) was obtained from an Indian Kala-azar patient and was maintained in golden hamsters (27). Amastigotes were obtained from infected hamster spleen and transformed into promastigotes. Promastigotes were maintained in M199 medium (Gibco) supplemented with 10% FBS (Gibco) and 1% Pen-Strep (Gibco) at 22°C. RAW264.7 cells were infected with stationary phase Ld promastigotes of the second to a fourth passage at a ratio of 1:10 (cell: Ld) for 6 hours (20).

### Optiprep density gradient centrifugation

For subcellular fractionation of organelles, a 3-30% continuous gradient was prepared using Optiprep density gradient medium of 60% iodixanol solution (Sigma-Aldrich) in a buffer composed of 78mM KCl, 4mM MgCl2, 8.4mM CaCl2, 10mM EGTA, 50mM HEPES (pH 7.0). Ice cold 1X PBS was used for washing before cells were lysed using a Dounce homogenizer (Kontes glass) in a buffer constituting 0.25M sucrose, 78mM KCl, 4mM MgCl2, 8.4mM CaCl2, 10mM EGTA, 50mM HEPES (pH 7.0) supplemented with 100μg/ml of cycloheximide, 0.5mM DTT and 1X PMSF. Cellular lysate was cleared by centrifuging twice at 1,000xg for 5 min before overlaying gently on top of the prepared continuous gradient. Ultracentrifugation was carried out for 5 hours at 133,000xg on a SW60-Ti (Beckman Coulter) rotor to resolve each gradient. After centrifugation, by aspirating from top, ten fractions were collected which were processed for analysis of proteins and RNA.

### Endosome enrichment by subcellular fractionation

Optiprep density gradient medium of 60% iodixanol solution (Sigma-Aldrich) was used to prepare a 3ml 5-10-15% step gradient (1ml of lighter solution overlaid gently on 1ml of heavier gradient) in a buffer containing 78mM KCl, 4mM MgCl2, 8.4mM CaCl2, 10mM EGTA, 50mM HEPES (pH 7.0) for separation of subcellular organelles. Cells were washed with 1X PBS and homogenized with a Dounce homogenizer (Kontes glass) in a buffer containing 0.25M sucrose, 78mM KCl, 4mM MgCl2, 8.4mM CaCl2, 10mM EGTA, 50mM HEPES (pH 7.0) supplemented with 100μg/ml of Cycloheximide, 0.5mM DTT and 1X PMSF. The lysate was clarified by centrifugation at 1,000xg for 5 min twice and 1ml of cleared lysate layered on top of the immediately prepared 3ml gradient at 4°C. The tubes were centrifuged at 133,000xg for 5 hours for separation of gradient and seven fractions of equal volume were collected by aspiration from the top for endosome/MVB isolation and subsequent analysis of proteins and RNA. For Endosome enrichment, top three fractions were pooled and diluted with a diluent buffer containing 78mM KCl, 4mM MgCl2, 8.4mM CaCl2, 10mM EGTA, 50mM HEPES (pH 7.0) to fill a 4ml tube and further ultracentrifuged at 133,000xg for another 2 hours. After centrifugation, endosome enriched membrane pellet was resuspended in a buffer containing 0.25M sucrose, 78mM KCl, 4mM MgCl2, 8.4mM CaCl2, 10mM EGTA, 50mM HEPES (pH 7.0) supplemented with 100μg/ml of cycloheximide, 0.5mM DTT and 1X PMSF, and this endosomal suspension was kept in ice before carrying out the in vitro assay.

### Cell-free in vitro reconstitution assay

The cell-free in vitro assays performed with endosomes were done following an earlier published protocol (28). Endosome enriched suspension isolated from either HEK293 or C6 cells were divided equally for experimental sets. As per the experiments described in the Results section, synthetic miRNA, and small RNA pool were specifically used as cargo in presence of ATP and incubation was carried out for 30 minutes at 37°C, followed by an Rnase protection assay as described in an earlier publication (14) and vesicles were washed and re-isolated by ultracentrifugation at 133,000 xg for 2 hours. Vesicle pellets were processed and analyzed for RNA and protein isolation.

### Immunoblotting

Western blotting of proteins was performed as mentioned in previous reports (28, 29). Protein samples from cellular lysate, subcellular fractionated samples, or immunoprecipitated proteins were subjected to SDS-polyacrylamide gel electrophoresis, followed by transfer of the same to PVDF nylon membrane overnight at 4°C. Membranes were blocked by using 3% BSA for 1 hour then specific required antibodies were used to probe the blot for at least 16h at 4°C. Membranes were incubated for 1h with horseradish peroxidase-conjugated secondary antibodies (1:8000 dilutions) at room temperature. Images of developed western blots were taken using an UVP BioImager 600 system coupled with VisionWorks Life Science software, version 6.8 (UVP).

### RNA Isolation and Real Time PCR

Total RNA was isolated by using TriZol or TriZol LS reagent (Invitrogen) according to the manufacturer’s protocol. Samples containing low amount of miRNA were precipitated by using isopropanol in presence of glycoblue co-precipitant (Thermofisher Scientific, USA). cDNA was prepared by taking 100-200ng of RNA from cellular RNA samples or by taking equal volume of RNA isolated from in vitro assay samples or EV samples by using a Taqman reverse transcription kit (Applied Biosystems). miRNA assays by real time PCR was performed using specific primers. Real time analyses by two-step RT-PCR were performed for quantification of miRNA levels on Bio-Rad CFX96TM real time system using Taqman (Applied Biosystems) based miRNA assay system. One third of the reverse transcription mix was subjected to PCR amplification with TaqMan Universal PCR Master Mix No AmpErase (Applied Biosystems) and the respective TaqMan reagents for target miRNA. Samples were analyzed in triplicates. For normalization of miRNA levels of cellular samples, U6 snRNA was used, whereas for in vitro assay samples done with synthetic miR-122, endosomal miR-146a was used for normalizing the level of miR-122. For each mRNA quantification, 200 ng of total cellular RNA was subjected to cDNA preparation followed by qPCR by the SYBR Green (Eurogentec) method. mRNA levels were normalized with GAPDH as the loading control. Each sample was analyzed in triplicates using the comparative Ct method.

### Cooperative miRNA association/dissociation kinetics assay

For the Cooperative miRNA association/dissociation kinetics assay, HEK293 cells were lysed by using lysis buffer (20 mM Tris-HCl pH 7.5, 150 mM KCl, 5 mM MgCl2 and 1 mM DTT) containing 0.5% Triton X-100, 0.5% sodium deoxycholate, 40 U/ml RNase inhibitor and 1X PMSF for 30 min at 4□C, followed by sonication of three pulses of 10 seconds each. Lysates were clarified by centrifuging at 16,000xg for 15 minutes. Protein G-agarose beads were blocked by using a solution of 5% BSA in 1X lysis buffer for 1 hour, then blocking buffer discarded and beads washed thrice using 1X IP buffer (20 mM Tris-HCl pH 7.5, 150 mM KCl, 5 mM MgCl2 and 1 mM DTT), followed by incubation with specific antibody solution in 1X lysis buffer (final dilution 1:100) for 3 hours at 4□C. The cleared cell lysates were incubated with primary antibody against HuR protein, bound with Protein-G Agarose bead (Invitrogen), and rotated overnight at 4□C. Thereafter, the beads were washed three times and resuspended in an Assay buffer containing 20mM Tris (pH 7.5), 5 mM MgCl2, 150mM KCl, 1X PMSF, 2mM DTT, and 40 U/ml RNase inhibitor. The synthetic miRNAs were added to the reaction mix as described in the experiments in the results section. The reaction was carried out at room temperature for 15 minutes with shaking. After the reaction, beads were washed three times and then divided into two equal parts, and each part was analyzed for bound proteins and RNAs by Western Blot and qRT-PCR, respectively.

### Small RNA pool isolation

Small RNA pool from cellular lysate was isolated using miRVANA miRNA isolation kit (Thermo fisher Scientific, USA) according to the manufacturer’s instructions. This small RNA pool was used as cargo for in vitro assay reactions, and the level of individual miRNA packaging was analyzed.

### Phosphate 5’ end-labeling of synthetic miRNA

50 pmoles of synthetic miR-122 RNA (22nt) was incubated with T4 Polynucleotide Kinase (PNK) enzyme (10 units/µl) and 1mM ATP in 1X T4 PNK buffer at 37°C for 30 minutes at static condition. Reaction was ended with Tris-EDTA solution and was filtered through mini Quick spin oligo column (Roche Diagnostics). RNA was extracted with TriZol LS and CHCl3 and precipitated with isopropanol at −20°C overnight in presence of glycoblue co-precipitant. RNA pellet was resupended in nuclease-free water to a final concentration of 1 pmoles/ µl.

### Preparation of recombinant HuR

Recombinant HuR protein was purified by procedure as previously described in detail (12). For the expression of HuR in E. coli, the construct consisting of HuR coding region in pET42a(+) was used. The protein was expressed in BL21DE3 E. coli cells codon Plus expression competent bacterial cells. IPTG-induced overnight cultures of E. coli BL21 were lysed by incubation with lysis buffer [20 mM Tris–HCl, pH 7.5, 300 mM KCl, 2 mM MgCl2, 5 mM β-mercaptoethanol, 50 mM imidazole, 0.5% Triton X-100, 5% glycerol, 0.5 mg/ml lysozyme, 1× EDTA-free protease inhibitor cocktail (Roche)] for 30 min followed by sonication (10 seconds, three pulses).The cleared lysate was incubated with pre-equilibrated Ni-NTA Agarose beads (Qiagen) for 4 h at 4°C. Bead washing was done using wash buffer (20 mM Tris–HCl, pH 7.5,150 mM KCl, 2 mM MgCl2, 5 mM β-mercaptoethanol, 50 mM imidazole, 0.5% Triton X-100, 5% glycerol) with rotation, on an end-to-end rotator, each for 10 min, at 4°C. His-tagged HuR protein was eluted by incubating beads with different elution buffers (20 mM Tris–HCl, pH 7.5,150 mM KCl, 2 mM MgCl2, 5 mM β-mercaptoethanol, 0.5% Triton X-100, 5% glycerol) of different and increasing imidazole concentrations, each for 15 min at 4°C with end-to-end rotation. Purified protein was kept in aliquots and stored at −80°C.

### Statistical Analysis

Software GraphPad Prism 5.00 (Graph Pad, San Diego) was used for analyzing the dataplots obtained from experiments performed at triplicate unless mentioned otherwise. Student’s t-test was done to determine P values. Significance was considered if P<0.05. Error bars represent mean±s.d.

## Data Availability Statement

All relevant data supporting the key findings of this study are available within the article and the Supplementary Information file. There is no Protein or RNA Sequencing data used in the manuscript to deposit in the repository domains.

## Acknowledgements

We thank Witold Filipowicz and Gunter Meister for the different constructs used in this study. We thank the Funding body, Dept. of Science and Technology (DST), Govt. of India along with the Council for Scientific and Industrial Research (CSIR), and University Grant Commission (UGC) for the SG. SNB was supported by The Swarnajayanti Fellowship (DST/SJF/LSA-03/2014-15) from Dept. of Science and Technology, Govt. of India. The work also received support from a High-Risk High Reward Grant (HRR/2016/000093) from Dept. of Science and Technology, Govt. of India, and CEFIPRA project grant 6003-J. SNB is currently supported by the Start-Up Support Grant of the University of Nebraska, USA, and the Lieberman Research Award, Department of Anesthesiology, UNMC support K.M.

## CRediT Author Contribution

Syamantak Ghosh: Data Curation, Formal Analysis, Investigation, Methodology. Kamalika Mukherjee : Data Curation, Formal Analysis, Investigation, Writing-Review and Editing, Conceptualization, Methodology, Supervision. Suvendra N Bhattacharyya : Data Curation, Formal Analysis, Investigation, Writing-Review and Editing, Conceptualization, Funding Acquisition, Methodology, Supervision, Writing-Review and Editing.

