## Supplementary material for "HuR-Dependent Cooperative Export of miRNAs From Activated Macrophages": Supple File

### Supplementary Figures with legends

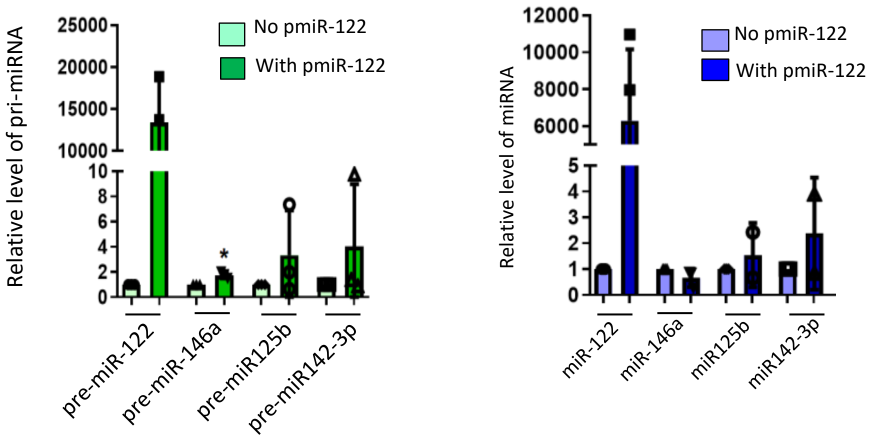


**Figure S1 Effect of coordinated miR-146a export by miR-122 on biogenesis of secondary miRNAs in RAW264.7 cells.**

Effect of cooperative miR-146a export on the mature and Pre-miRNA levels of secondary miRNAs known to be regulated by miR-146a. Pre-miRNAs and mature miRNAs were quantified by qRT-PCR, and the 0ng pmiR-122 plasmid transfection condition dataset was taken as a unit (n=3). For pmiR122 set, 1000ng plasmids per $8x10^6$ cells were used for transfection.

Data information: In all the experimental data, error bars are represented as mean with SD. Relative levels of miRNAs were normalized with U6 snRNA level by $2^{-\Delta\Delta Ct}$ method.

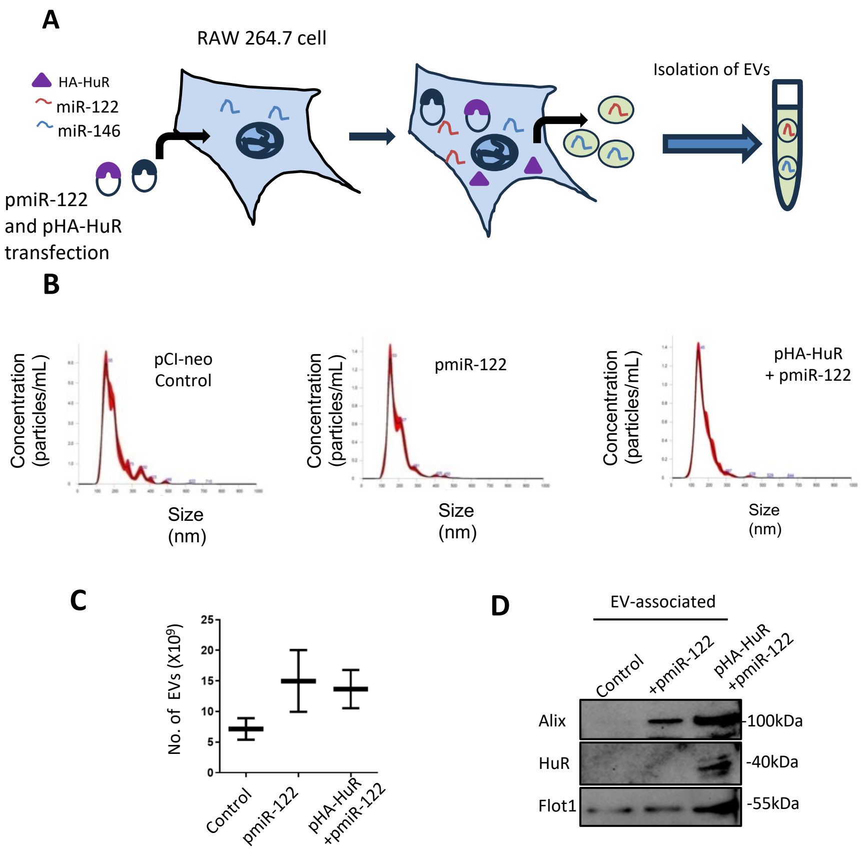


**Figure S2 Ectopic expression of miR-122 and HA-HuR enhances the EV-production in RAW264.7 cells.**

**A.** The miR-122 expression vector (pmiR-122) and HA-HuR expression vector (pHA-HuR) were transfected separately and together in RAW.264.7 cells. Cells secreted EVs packed with ectopically expressed miR-122 and endogenously expressed miR-146a were quantified. pCI-neo was used as the control plasmid for transfection. The conditioned media were used to isolate EVs.

**B.** Nanoparticle tracking analysis (NTA) graphs of EVs from the sample sets of control, miR-122 expressing or miR-122 and HA-HuR co-expressing cells.

**C.** Analysis of the number of exosomes estimated by NTA (n=2).

**D.** Analysis of proteins associated with EVs by western blots analysis from the experiment performed in panel A.

Data information: In all the experimental data, error bars are represented as mean with SD. Positions of molecular weight markers are marked and shown with the respective Western blots.

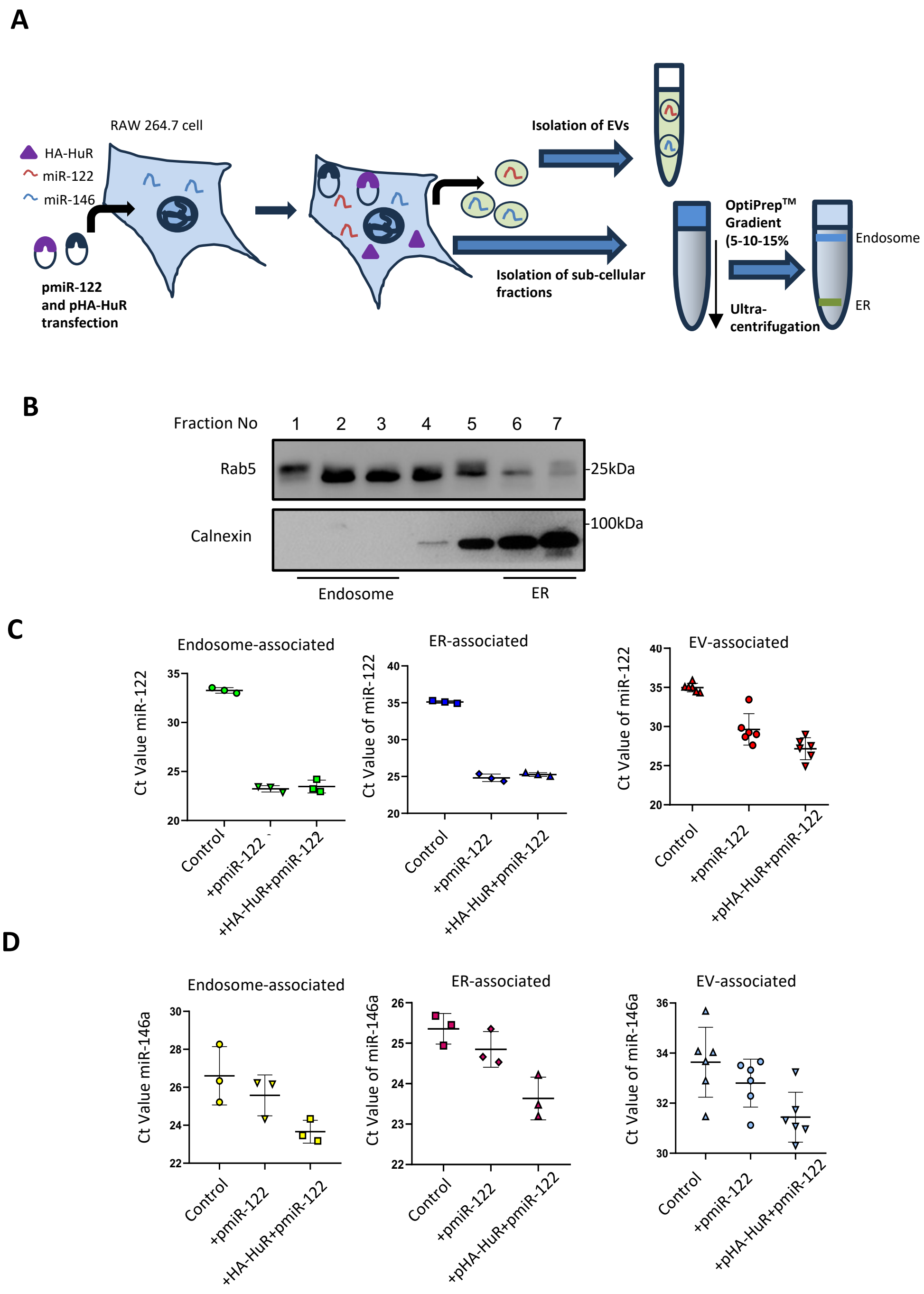


**Figure S3 Effect of miR-122 and HA-HuR expression on the subcellular and EV-associated miR-146a levels**

**A.** The miR-122 and HA-HuR expression vectors were expressed separately and together in RAW.264.7 cells. Secreted EVs packed with ectopically expressed miR-122 and endogenously expressed miR-146a were collected from the conditioned media. The lysates of transfected cells were subjected to fractionation on 5-10-15% OptiPrep$^{TM}$ density gradient centrifugation to separate endosomes (Fractions 1-3) and Endoplasmic Reticulum (ER, Fractions 6-7). These subcellular organelles and vesicles were analyzed for the change in EV miRNA secretion level and the change in the subcellular localization of miRNAs due to the cooperativity of miRNA packaging.

**B.** Western blots of endosomal marker Rab5 and ER marker Calnexin from the fractions obtained from the density gradient centrifugation of RAW264.7 cell lysates.

**C.** Cells expressing control pCI-neo plasmid, or p-miR-122 alone or with pHA-HuR were harvested to analyze miRNA levels from various vesicular compartments to observe the changes in the localization of miRNA across subcellular compartments. Subcellular fractionation of cell lysates was done, and endosomal (left panel) and ER (center panel) fractions were used to isolate the RNA. EVs (right panel) were isolated from cell culture-conditioned media. QRT-PCR quantified the miR-122 level, plotting Ct values (n=3). Lower Ct value signifies higher miRNA abundance.

**D.** QRT-PCR also analyzed the level of miR-146a from the same experimental samples as in C (n=3).

Data information: In all the experimental data, error bars are represented as mean with SD. Positions of molecular weight markers are marked and shown with the respective Western blots.

**Supplementary Table 1: List of Plasmids and Chemicals**

| Name | Reference/Source | Description |
|---|---|---|
| pmiR-122 | As described by Chang J. | Plasmid encoding pre-miR-122 under a constitutive U6 promoter |
| pHA-HuR | Kind gift of Witold Filipowicz | Full length HuR with HA coding sequence in pCI-Neo vector |
| GW4869 | Calbiochem, CA | Exosome secretion blocker, 10μM Conc. |
| LPS | Calbiochem, CA | *E. coli* O111:B4 Lipopolysaccharide |

**Supplementary Table 2: Details of miRNA primers used for Taqman based quantification**

| Name | Assay ID |
|---|---|
| miR-122 | 000445 |
| miR-146a | 000468 |
| miR-155 | 002571 |
| let-7a | 000377 |
| miR-21 | 000397 |
| miR-125b | 000449 |
| miR-142-3p | 000464 |
| U6 SnRNA | 001973 |

**Supplementary Table 3: Details of Antibodies used for western blot (WB), Immunofluorescence (IF) and immunoprecipitation (IP)**

| Antibody Name | Raised in | Dilution | Source | Catalogue No. |
|---|---|---|---|---|
| p38 | Rabbit | WB- 1:1000 | Cell Signaling | #9212 |
| P-p38 | Rabbit | WB- 1:1000 | Cell Signaling | #9211 |
| ERK | Rabbit | WB- 1:1000 | Cell Signaling | #9102 |
| P-ERK | Rabbit | WB- 1:1000 | Cell Signaling | #9101 |
| c-PARP | Rabbit | WB- 1:1000 | Cell Signaling | #5625 |
| Alix | Mouse | WB-1:100 | Santa Cruz | sc53538 |
| Calnexin | Rabbit | WB- 1:10,000 | Bethyl | A303-696A |
| Rab5a | Rabbit | WB-1:1000<br>IF- 1:100 | Cell Signaling | #3547S |
| Rab7a | Rabbit | WB-1:1000 | Cell Signaling | #9367S |
| EEA1 | Rabbit | WB-1:1000 | Cell Signaling | #2411S |
| HuR | Mouse | WB-1:1000 | Santa Cruz | sc-5261 (3A2) |
| HA | Rat | WB-1:1000<br>IP- 1:100<br>IF:100 | Roche | 11867431001 |
| β-Actin | Mouse monoclonal (HRP-conjugated) | WB- 1:10,000 | Sigma Aldrich | A3854 |
| Flotilin-1 | Rabbit | WB-1:1000 | Cell Signaling | #18634S |
| Ago2 (eIF2C2) | Mouse | WB-1:500 | Abnova | H00027161-M01 |
| GM130 | Rabbit | WB-1:1000 | Abcam | EP892Y |

**Supplementary Table 4: List of Synthetic single stranded RNA and anti-miR oligonucleotides**

| Name | Sequence/Catalogue No. |
|---|---|
| miR-122-5p | 5'- UGGAGUGUGACAAUGGUGUUUG-3' |
| miR-146a-5p | 5'-UGAGAACUGAAUUCCAUGGGUU-3' |
| miR-155-5p | 5'-UUAAUGCUAAUCGUGAUAGGGGUU-3' |
| Anti-miR-146a-5p | Ambion (AM10722) |
| Anti-miR Negative Control | Ambion (AM17010) |

**Supplementary Table 5: Details of mRNA primers used for SYBR-Green based quantification**

| Target | 5' Forward Primer 3' | 5' Reverse Primer 3' |
|---|---|---|
| TNF-α | GTCTCAGCCTCTTCTCATTCC | TCCACTTGGTGGTTTGCTA |
| IL-1β | GACCTTCCAGGATGAGGACA | CCTTGTACAAAGCTCATGGAG |
| IL-10 | TGCTAACCGACTCCTTAATGC | ATCACTCTTCACCTGCTCCAC |
| IL-6 | AGGATACCACTCCCAACAGA | GTACTCCAGAAGACCAGAGGA |
| Amastin | GGGGTTCAAAGTTCGAGTGC | AGCAAAGAGCAGCAGCACAG |
| GAPDH | CAGGGGGGAGCCAAAAGGG | CTTGGCCAGGGGTGCTAAGC |
